# Polyelectrolyte brush bilayers at thermal equilibrium: Density functional theory and molecular dynamic simulations

**DOI:** 10.1101/2022.02.24.481861

**Authors:** Mike J. Edwards

## Abstract

In this article, the equilibrium properties of Polyelectrolyte brush bilayers is studied. The density functional theory framework as well as the benchmark molecular dynamic simulations are employed to indicate the correct equation of state. Both techniques turn out that the equation of state neutral polymer brush bilayers scales with the wall distance as *D*^−4^. Additionally, it turns out that the electrostatic interactions does not change the universal power laws. The only difference is that the charged systems posses larger pressure due to stored electrostatic energy.

**SIGNIFICANCE:** Studying Polyelectrolyte brush bilayers is important because it exists widely in nature, biology and industry. In biology, they could be found in mammalian synivial joints as well as in the cellular structures.

## INTRODUCTION

Large linear macromolecular structures are polymers which consist of repeating units or monomers (1, 5, 20). The thermal fluctuations arising from the random kicks by solvent molecules cause monomers to fluctuate. The fluctuations of monomers in long time scales make polymer chain to maintain its average coil size (1, 5, 20). The average coil size of a chain in absence of the excluded volume interactions scales as *∼N*^1/2^ and in presence of the excluded volume interactions scales as *∼N*^3/5^. When the polymer chains are densely grafted to a surface, the steric repulsion between the nearby chains stretches the chains in perpendicular direction and a polymer brush forms (2, 7, 8, 11). The average brush height scales as ∼ *N* (7–11, 24–26). Polymer brush bilayers (P.B.B.) form when two opposing brush covered surfaces compressed upon each other in the way that brushes interpenetrate. In this condition, the brush height scales as ∼ *N* but it does not follow a power law by the wall distance (27). It has been recently turned out that the equation of state of a P.B.B. scales as (*N* /*D*)^4^ with *D* the wall distance (27). Note that, there has been studies which propose different power laws for P.B.B. (12–15) which are calculated based on blob picture. The P.B.B. is experimentally studied to show that it has fascinating lubrication properties and nature uses it in mammalian synivial joint (16–18). The Polyelectrolyte brush bilayers (P.E.B.B.) forms when the monomer carry charge. Some experimental studies of the P.E.B.B. proposes that they have better lubrication properties rather than neutral ones (12–18). However, recently it has been theoretically shown that the charge effect has no role in lubrication (29). In this article, I solely deal with the problem of the P.E.B.B. at thermal equilibrium. I challenge the theoretical results already published with respect to power laws and the way they approach the charge effect. I start approaching this problem through two independent techniques D.F.T. and M.D. and get good agreement.

### DENSITY FUNCTIONAL THEORY FRAMEWORK

The D.F.T. framework is the best theoretical tool to tackle the many body systems. To apply it to the P.E.B.B. problem, I use the same approach proposed in (6, 19) for polymer brushes and I extend it to the P.E.B.B.. In this technique, the grand potential functional of the system is build by balancing entropic elasticity and the excluded volume interactions. The charge effect is included in the second Virial coefficient. After minimizing the grand potential functional with respect to density, a set of coupled equations must be solved. By putting the results of that back into the grand potential, the desired result is obtained. To calculate the equation of state or pressure, the Gibbs-Duhem relation is used (3). The details of calculations could be found in (27). The resulting equation of state is obtained after that procedure,

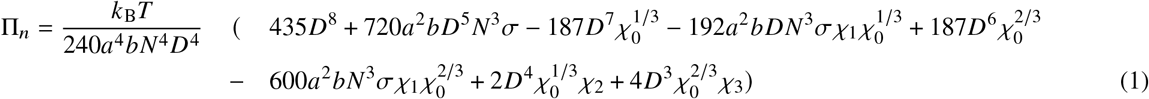

where, *a* is the Kuhn length or monomer size, *b* the second Virial coefficient, *N* the degree of polymerization, *D* the wall distance and *σ* the grafting density. Also, the following variables are introduced to simplify the expression,

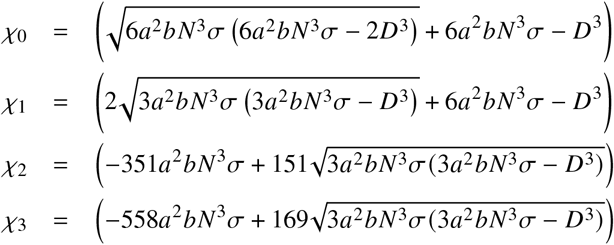

Fitting Eq. (1) reveals that it scales with system parameters as following,

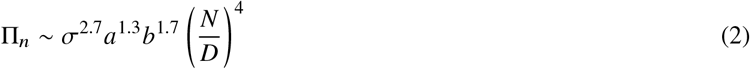

### MOLECULAR DYNAMIC SIMULATIONS

The most reliable numerical technique for studying polymer solutions is the molecular dynamic simulations (M.D.). Here, I utilize the coarse-grained M.D. simulations proposed by Kremmer and Grest known as KG model (22). In this model, the monomers are represented by the Lennard-Jones potential as follows,

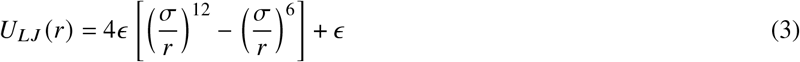

with *∈* the energy scale and the *σ* the monomers size. Note that this potential is cutt-off at *r* = 2^1/6^*σ* to simulate good solvent conditions. The connectivity along the backbone of the chains is intoduced by Finitely Extensible Nonlinear Elastic potential (F.E.N.E.) as follows,

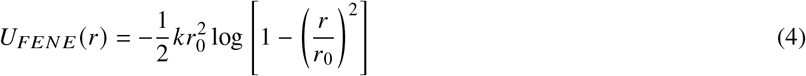

with *k* the elastic constant and *r*_0_ the maximum allowed extension. To include the electrostatic interactions, I use Debye-Hückel potential as follows,

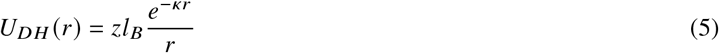

where the Bjerrum length which is the length in which thermal and electrostatic energies balance, is defined as follwing,

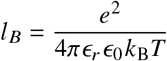

and the Debye screening length *κ*^−1^ is defined as following,

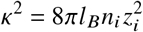

And finally, the thermal fluctuations is implemented through Langevin thermostat which is given as a random force and a dissipative force,

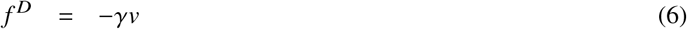

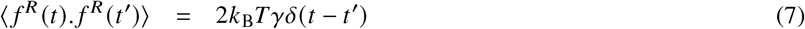

with *ϒ* the monomer friction coefficient and *υ* the monomer velocity. The Newton’s equations of motion are discretized through Leap-Frog algorithm (21) and the simulation parameters are listed in Table (1).

**Table 1:** The numerical values of the system parameters

|  |  |  |
| --- | --- | --- |
| Temperature | $T$ | $1.68 \epsilon / k_B$ |
| MD simulations time step | $\Delta t$ | $2 \times 10^{-3} (m\sigma^2/\epsilon)^{1/2}$ |
| MD simulations run time | $t$ | $10^4 \Delta t$ |
| Lennard-Jones cutoff for good solvent | $r_c$ | $2^{1/6} \sigma$ |
| FENE maximum extension | $r_0$ | $1.5 \sigma$ |
| FENE elastic constant | $k$ | $30 \epsilon / \sigma^2$ |
| Hydrodynamic friction coefficient | $\gamma$ | $5 (m\epsilon/\sigma^2)^{1/2}$ |
| Bjerrum length | $l_B$ | $0.7 \sigma$ |
| CI concentration | $n_i$ | $10^{-1} \sigma^{-3}$ |
| CI valency | $z_i$ | 1 |
| Monomer valency | $z$ | 1 |
| Debye screening length for $l_B = 0.7 \sigma$ | $\kappa^{-1}$ | $0.75 \sigma$ |
| Kuhn length | $a$ | $0.96 \sigma$ |
| Second Virial coefficient for polymer in good solvent | $b$ | $2.08 \sigma^3$ |
| Second Virial coefficient for P.E.B.B. in good solvent $l_B = 0.7$ | $b$ | $2.99 \sigma^3$ |
| Squeeze velocity | $v_s$ | $10(\epsilon/m)^{1/2}$ |

The simulations are performed in two parts. First, the brushes start squeezing from their initial distance by high relative velocity of 10*σ* and they stop in desired distance. In the second part, the walls stay in that distance for a long time and relax. In the second part the pressure measurement is performed. To calculate pressure in terms of the degree of polymerization, the walls squeeze to *D* = 8*σ* and the systems with different *N* are measured. The results of both technique are represented in Fig. (1) where the pressure is shown in terms of wall distance as well as degree of polymerization. Both graphs show a good agreement between the D.F.T. framework and M.D. simulations. They support the idea that the equation of state scales as (*N*/*D*)^4^ and also the fact that the electrostatic interactions do not change the power law but they offer a larger pressure.

**Figure 1:**
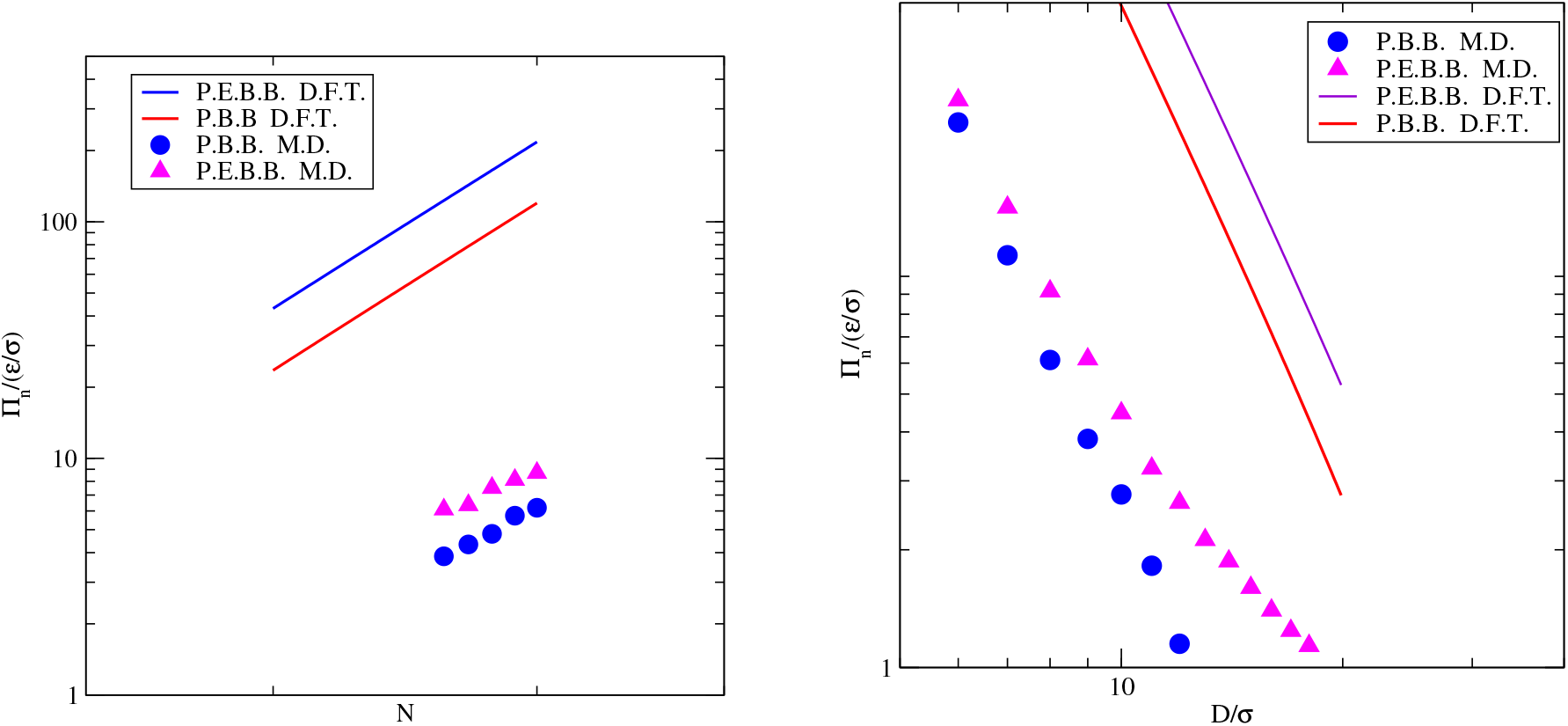
Left: The equation of state in terms of the degree of polymerization. Right: The equation of state in terms of the wall distance. In both graphs the solid lines show the DFT results and the symbols show the MD simulations results.

### CONCLUDING REMARKS

In the present article, the problem of P.E.B.B. at thermal equilibrium is studied through D.F.T framework as well as M.D. simulations. It turns out that the pressure of the system scales as ∼ (*N* / *D*)^4^ and the previous studies which propose ∼ *D*^−7/2^ and ∼ *N* need to be reviewed. In particular, the work already published (15). In the context of the charge effect, my study shows that charge effect does not change the power laws. In contrast to the results of the work already published which suggests that P.E.B.B. behave like a melt. Since, in my study I use two independent techniques and both of them are highly reproducible, I would conclude that the blob picture is not a reliable technique to approach many body systems like polymers. One of weaknesses of the blob picture is that it hypothetically considers a blob in which the excluded volume interactions occur and beyond that blob there is no interactions. Something similar to the electrostatic interactions. But the point is that the excluded volume interactions are short range and the screening length does not exist for them. Another weakness of the blob picture is its simplicity and the fact that it can not capture the complexity of systems like polymer solutions. In the work already published, one could find many perfect agreement between blob picture and M.D.. But as I am well familiar with how research has been done in that work, I would say that the author may matched its theory up with the simulation results. This is not how research in science works because to prove the correctness of a result, one should find agreement between completely independent techniques. At the end, the results of this study may be useful in our understanding of P.E.B.B. and its biological applications such as artificial synovial joints.

## ACKNOWLEDGMENTS

I thank Cold Spring Harbor Laboratory (CSHL) and Regeneron Pharmaceuticals Inc. for financial support of this study and I thank Wolfram Research for providing Mathematica software, the Weizmann institute of science for the XMGrace software, the NIH for VMD software, the Redhat for Fedora linux operating system and the Overleaf latex editor.

